# GTDB-Tk v2: memory friendly classification with the Genome Taxonomy Database

**DOI:** 10.1101/2022.07.11.499641

**Authors:** Pierre-Alain Chaumeil, Aaron J. Mussig, Philip Hugenholtz, Donovan H. Parks

## Abstract

The Genome Taxonomy Database (GTDB) and associated taxonomic classification toolkit (GTDB-Tk) have been widely adopted by the microbiology community. However, the growing size of the GTDB bacterial reference tree has resulted in GTDB-Tk requiring substantial amounts of memory (~320 GB) which limits its adoption and ease of use. Here we present an update to GTDB-Tk that uses a divide-and-conquer approach where user genomes are initially placed into a bacterial reference tree with family-level representatives followed by placement into an appropriate class-level subtree comprising species representatives. This substantially reduces the memory requirements of GTDB-Tk while having minimal impact on classification.

**Availability:** GTDB-Tk is implemented in Python and licenced under the GNU General Public Licence v3.0. Source code and documentation are available at: https://github.com/ecogenomics/gtdbtk.

## 1 Introduction

GTDB-Tk has been used to assign taxonomic classifications to tens of thousands of bacterial and archaeal isolate genomes and meta-genome-assemble genomes (MAGs) recovered from environmental and human-associated samples (Chaumeil et al., 2019; Almeida et al., 2021; Nayfach et al., 2021). These classifications are consistent with the GTDB framework and based on the same relative evolutionary divergence (RED) and average nucleotide identity (ANI) criteria for circumscribing taxa (Parks et al., 2019 and 2022). A primary step in assigning classifications is placing genomes into the GTDB bacterial or archaeal reference trees using the maximum-likelihood (ML) placement tool pplacer (Matsen et al. 2010). Unfortunately, ML placement with pplacer is a memory intensive operation requiring ~320 GB of RAM when using the GTDB R07-RS207 bacterial reference tree comprised of 62,291 genomes. Adding to this challenge is the lack of favourable alternatives to pplacer as the EPA-ng ML placement method requires more memory than pplacer and distance-based methods such as APPLES-2 have inferior performance (Barbera et al., 2019; Balaban et al., 2021; Koning et al., 2021). The GTDB bacterial reference tree has been growing rapidly in size with each GTDB release (Parks et al., 2022) which has ultimately made the memory requirements of GTDB-Tk impractical. Here we show that the memory requirements of GTDB-Tk are reduced by dividing the GTDB bacterial reference tree into class-level subtrees and demonstrate that taxonomic classifications are largely unimpacted by this change.

## 2 Methods

GTDB-Tk v2 divides the GTDB bacterial reference tree into class-level subtrees to reduce memory requirements. Placement of a genome with pplacer now consists of two steps. First, a genome is placed into a backbone tree consisting of a single genome representative for each family (see *Supp. Methods*). If the genome is assigned to a class within this backbone tree, it is then placed into a class-level subtree to obtain a more refined placement for the genome. The class-level subtrees were constructed in a greedy manner with the maximum size of a class-level subtree being set based on the number of species representatives in the largest class (*Gammaproteobacteria* with 9,582 genomes in GTDB R07-RS207). Each class-level subtree was formed by selecting the largest class in the reference tree and traversing towards the root until the subtree contained at most 10,540 genomes (10% more than the *Gammaproteobacteria*). This subtree was then pruned from the reference tree and the procedure repeated until all classes were assigned to a class-level subtree. For GTDB R07-RS207, this results in 7 class-level trees. Each class-level subtree was then expanded to contain a single genome from each phylum in order to allow query genomes to be placed as the most basal member of a class.

Final taxonomic classifications use the same RED and ANI criterion as GTDB-Tk v1 (Chaumeil et al., 2020) with the following additional rules:

1. A genome not placed into a class-level subtree is assigned the classification determined in the backbone tree.
2. A genome placed into a class-level subtree and assigned to a phylum belonging to one of the classes contained in the subtree is assigned the classification determined in the subtree.
3. Otherwise, the genome is classified by taking the lowest common ancestor between the backbone and class-level subtree.

## 3 Results

Here we demonstrate that the taxonomic classifications produced by the divide-and-conquer approach implemented in GTDB-Tk v2 are nearly equivalent to those produced by GTDB v1 while providing a substantial reduction in required memory.

### 3.1 Similarity of classifications on diverse sets of genomes

The concordance between GTDB-Tk v1 and v2 classifications was first assessed using 16,710 bacterial genomes from the GEMs dataset (Nayfach et al., 2021) that represent novel taxa relative to GTDB R07-RS207 (**Table 1**). Only 12 genomes (0.07%) did not have identical classifications between GTDB-Tk v1 and the divide-and-conquer approach used in GTDB-Tk v2 (**Supp. Table 1**). The majority of incongruence was due to genomes being over- (6 genomes) or under-classified (4 genomes) by a single taxonomic rank. Only 2 genomes had conflicting taxonomic assignments, and these were both relatively poor-quality genomes assigned as new classes in alternative phyla (**Supp. Table 1**).

**Table 1.** Novelty of GEM genomes relative to GTDB R07-RS207 based on GTDB-Tk v1 classifications.

| *Taxon Novelty* | *No. genomes* | *GTDB-Tk v2 classifications relative to GTDB-Tk v1 classifications* |  |  |  |
| --- | --- | --- | --- | --- | --- |
|  |  | *Congruent* | *Conflict* | *Underclassified* | *Overclassified* |
| Novel phylum | 3 | 2 | 0 | 0 | 1 |
| Novel class | 42 | 36 | 2 | 2 | 2 |
| Novel order | 144 | 143 | 0 | 0 | 1 |
| Novel family | 543 | 540 | 0 | 1 | 2 |
| Novel genus | 3,222 | 3,221 | 0 | 1 | 0 |
| Novel species | 12,756 | 12,756 | 0 | 0 | 0 |

GTDB-Tk v1 and v2 classifications were further evaluated by dereplicating the ~60,000 genomes introduced in GTDB R07-RS207 to 23,548 genomes by randomly selecting a single genome per species. These 23,548 genomes were then classified using the GTDB-Tk R06-RS202 reference package to further evaluate classifications on genomes with varying degrees of taxonomic novelty and to ensure results are robust with different GTDB reference packages (**Supp. Table 2**). Only 13 genomes (0.06%) had different GTDB-Tk v1 and GTDB-Tk v2 classifications with 5 having conflicting assignments, 5 being overclassified, and 3 being underclassified (**Supp. Table 3**).

### 3.2 Reduced memory requirements

The divide-and-conquer approach implemented in GTDB-Tk v2 reduces the maximum memory requirements from ~320 GB to <55 GB when run with the GTDB R07-RS207 reference trees. GTDB-Tk v2 also runs 22% to 35% faster when processing 1,000 genomes with 1 to 64 CPUs (**Supp**. **Fig. 1A**) and is >40% faster when processing 5,000 genomes using 32 CPUs (**Supp. Fig. 1B**).

## 4 Summary

GTDB-Tk v2 requires only a sixth of the memory of GTDB-Tk v1 while providing almost identical classifications. More importantly, the divide-and-conquer approach used in GTDB-Tk v2 allows memory requirements to be controlled by tailoring the size of the largest subtree. This ensures GTDB-Tk can continue to be used on readily available computing hardware even as that size of the GTDB bacterial reference tree increases.

## Acknowledgements

We thank Morgan Price for insightful advice on using FastTree with pplacer, Brian Kemish for his help in maintaining our computing infrastructure, Maria Chuvochina and Christian Rinke for helpful discussions, and the GTDB-Tk community for their bug reports and suggestions on features to improve GTDB-Tk.

