## Supplementary material for "GTDB-Tk v2: memory friendly classification with the Genome Taxonomy Database"

**Supplementary Methods**

The backbone tree used in GTBD-Tk v2 consists of a single genome for each family. Assembly quality and NCBI metadata was used to establish the genome selected to represent each family. Specifically, each genome was assigned a score as follows:

- CheckM completeness estimate – 5 × CheckM contamination estimate
- -5 × number of contigs / 100
- -5 × number of ambiguous bases / 10000
- -5 × percentage of gaps in GTDB multiple sequence alignment
- -100 if assembly is from a MAG or SAG

The highest scoring genome in each GTDB family was selected for use in the GTDB-Tk v2 backbone tree.

**Supp. Table 1.** Incongruent GTDB-Tk v1 and v2 classifications across 16,710 GEM MAGs.

| *GEMs ID* | *CheckM completeness (%)* | *CheckM contamination (%)* | *GTDB-Tk v1 classification* | *GTDB-Tk v2 classification* | *Type of Change* |
|---|---|---|---|---|---|
| 3300017989_37 | 75.27 | 0 | d__Bacteria; p__4572-55; c__ | d__Bacteria; p__Gemmatimonadota; c__ | Conflict |
| 3300024433_39 | 62.46 | 0 | d__Bacteria; p__TA06_A; c__ | d__Bacteria; p__Gemmatimonadota; c__ | Conflict |
| 3300025546_16 | 96.27 | 0 | d__Bacteria; p__ | d__Bacteria; p__Aerophobota; c__ | Overclassified |
| 3300005529_81 | 70.3 | 0 | d__Bacteria; p__Chloroflexota; c__ | d__Bacteria; p__Chloroflexota; c__Ktedonobacteria; o__ | Overclassified |
| 3300013133_46 | 89.51 | 1.12 | d__Bacteria; p__Firestonebacteria; c__ | d__Bacteria; p__Firestonebacteria; c__D2-FULL-39-29; o__ | Overclassified |
| 3300025847_116 | 64.12 | 0 | d__Bacteria; p__Caldisericota; c__Caldisericia; o__JAAYUI01; f__ | d__Bacteria; p__Caldisericota; c__Caldisericia; o__JAAYUI01; f__JAAYUI01; g__ | Overclassified |
| 3300025856_28 | 56.8 | 1.23 | d__Bacteria; p__OLB16; c__OLB16; o__ | d__Bacteria; p__OLB16; c__OLB16; o__SURF-12; f__ | Overclassified |
| 3300027863_103 | 83.62 | 3.45 | d__Bacteria; p__Actinobacteriota; c__Thermoleophilia; o__UBA2241; f__ | d__Bacteria; p__Actinobacteriota; c__Thermoleophilia; o__UBA2241; f__UBA2241; g__ | Overclassified |
| 3300000227_24 | 93.28 | 0.84 | d__Bacteria; p__Desulfobacterota_D; c__ | d__Bacteria; p__ | Underclassified |
| 3300006083_11 | 81.72 | 0.81 | d__Bacteria; p__Aquificota; c__Desulfurobacteriia; o__Desulfurobacteriales; f__Desulfurobacteriaceae; g__ | d__Bacteria; p__Aquificota; c__Desulfurobacteriia; o__Desulfurobacteriales; f__ | Underclassified |
| 3300026127_6 | 81.7 | 4.46 | d__Bacteria; p__Desulfobacterota_D; c__UBA1144; o__RKRQ01; f__ | d__Bacteria; p__Desulfobacterota_D; c__UBA1144; o__ | Underclassified |
| 3300026488_14 | 68.64 | 0 | d__Bacteria; p__Muirbacteria; c__ | d__Bacteria; p__ | Underclassified |

**Supp. Table 2.** Novelty of 23,548 GTDB R07-RS207 genomes relative to R06-RS202 based on GTDB-Tk v1 classifications.

| | *No. genomes* | *GTDB-Tk v2 classifications relative to GTDB-Tk v1 classifications* |
|---|---|---|
| Novel phylum | 3 | 3 congruent |
| Novel class | 32 | 28 congruent; 3 conflict; 1 underclassified |
| Novel order | 199 | 194 congruent; 2 conflict; 1 underclassified; 2 overclassified |
| Novel family | 580 | 577 congruent; 1 underclassified; 2 overclassified |
| Novel genus | 3,393 | 3,392 congruent; 1 overclassified |
| Novel species | 12,616 | 12,616 congruent |
| Known Species | 6,725 | 6,725 congruent |

**Supp. Table 3.** Incongruent GTDB-Tk v1 and v2 classifications across 23,548 MAGs introduced in GTDB R07-RS207.

| *GEMs ID* | *CheckM completeness (%)* | *CheckM contamination (%)* | *GTDB-Tk v1 classification* | *GTDB-Tk v2 classification* | *Type of Change* |
|---|---|---|---|---|---|
| GB_GCA_018818825.1 | 97.75 | 1.12 | d__Bacteria; p__UBA9089; c__UBA9089; o__ | d__Bacteria; p__UBA9089; c__CG2-30-40-21; o__ | Conflict |
| GB_GCA_016209155.1 | 86.41 | 4.03 | d__Bacteria; p__Aerophobota; c__ | d__Bacteria; p__UBP18; c__UBA7526; o__ | Conflict |
| GB_GCA_016926495.1 | 76.57 | 1.68 | d__Bacteria; p__Bdellovibrionota; c__FAC87; o__ | d__Bacteria; p__Bdellovibrionota; c__YA12-FULL-61-11; o__ | Conflict |
| GB_GCA_016190245.1 | 75.18 | 0.84 | d__Bacteria; p__J088; c__ | d__Bacteria; p__SpSt-318; c__ | Conflict |
| GB_GCA_016783345.1 | 63.18 | 0 | d__Bacteria; p__NPL-UPA2; c__ | d__Bacteria; p__CAIJMQ01; c__ | Conflict |
| GB_GCA_017861215.1 | 81.27 | 0.81 | d__Bacteria; p__Desulfobacterota_F; c__Desulfuromonadia; o__Geobacterales; f__Geobacteraceae; g__ | d__Bacteria; p__Desulfobacterota_F; c__Desulfuromonadia; o__Geobacterales; f__Geobacteraceae; g__VEOV01; s__ | Overclassified |
| GB_GCA_018829865.1 | 70.89 | 2.97 | d__Bacteria; p__Patescibacteria; c__Gracilibacteria; o__UBA1369; f__ | d__Bacteria; p__Patescibacteria; c__Gracilibacteria; o__UBA1369; f__PNNO01; g__ | Overclassified |
| GB_GCA_016927475.1 | 90.37 | 2.56 | d__Bacteria; p__Acidobacteriota; c__Aminicenantia; o__Aminicenantales; f__ | d__Bacteria; p__Acidobacteriota; c__Aminicenantia; o__Aminicenantales; f__Aminicenantaceae; g__ | Overclassified |
| GB_GCA_015231965.1 | 95.66 | 0 | d__Bacteria; p__Proteobacteria; c__Magnetococcia; o__ | d__Bacteria; p__Proteobacteria; c__Magnetococcia; o__Magnetococcales; f__ | Overclassified |
| GB_GCA_016212085.1 | 99.49 | 0 | d__Bacteria; p__Nitrospirota; c__Nitrospiria; o__ | d__Bacteria; p__Nitrospirota; c__Nitrospiria; o__SBBL01; f__ | Overclassified |
| GB_GCA_016180645.1 | 89.33 | 3.36 | d__Bacteria; p__SpSt-318; c__; o__; f__; g__; s__ | d__Bacteria; p__ | Underclassified |
| GB_GCA_015231815.1 | 79.81 | 1.71 | d__Bacteria; p__Nitrospinota; c__UBA7883; o__ | d__Bacteria; p__ | Underclassified |
| GB_GCA_015233785.1 | 61.19 | 2.1 | d__Bacteria; p__Proteobacteria; c__Magnetococcia; o__Magnetococcales; f__ | d__Bacteria; p__Proteobacteria; c__Magnetococcia; o__ | Underclassified |

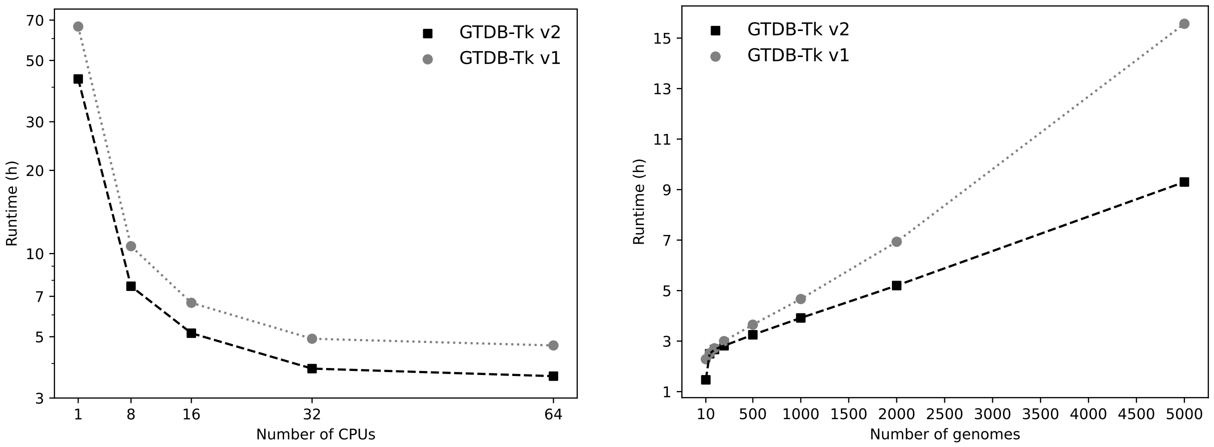


**Supp. Figure 1**. Reduced processing time of GTDB-Tk v2 compared to GTDB-Tk v1. (**a**; **left**) Processing time (log scale) for 1,000 randomly selected GEM MAGs for increasing numbers of CPUs. (**b**; **right**) Processing time with 32 CPUs on increasing numbers of randomly selected GEM MAGs. Tests were run on a machine with 4 AMD EPYC 7402 24-Core Processor and 512 GB of RAM.
